# Host species composition, not priority effects, determines infection risk in multihost-multiparasite amphibian communities

**DOI:** 10.1101/2025.06.18.660433

**Authors:** David R Daversa, Andy Fenton, Lola M Brookes, Goncalo Rosa, Chris Sergeant, Trenton WJ Garner

## Abstract

Novel parasite emergences are posing uncertain threats to wildlife communities worldwide. Because parasites typically infect multiple host species and interact with other parasites, the degree of threat posed by these emergences is known to depend both on a community’s host species composition and history of exposure to other parasites. What is not known is how these two factors – multi-host effects and multi-parasite effects – interact to shape the outcomes of parasite emergences. We experimentally investigated the outcomes of sequential emergences by two amphibian parasites of major concern to global biodiversity, *Batrachochytrium dendrobatidis* (*Bd*) and Ranavirus (*Rv*). Mimicking sequential *Bd* and *Rv* emergence in communities comprising either larval common toads (*Bufo bufo*), larval common frogs (*Rana temporaria*), or both host species, showed that host community composition influenced subsequent patterns of *Bd* and *Rv* infection much more than the presence of the other parasite. *Bd* prevalence varied predictably with host community composition; toads exhibited consistently higher *Bd* infection probabilities than frogs across all scenarios. Variation in community-level prevalence therefore was explained by differences in the relative abundance of toads. In contrast, community-level *Rv* prevalence was not consistently driven by a single host species. Going from single-host to two-host communities reversed relative *Rv* infection probabilities in frogs versus toads; when kept separately, frogs had a higher *Rv* infection probability than toads, but toads had a higher infection probability than frogs when the two species were co-housed. Host species composition can therefore alter multi-parasite infection dynamics not only through changes in the abundance of primary host species, but also through changes in which species appear to act as the primary hosts. These insights suggest that *Bd*-*Rv* dynamics in single-host scenarios may not reflect those occurring in natural multihost communities, and the observed context-dependency in the apparent primary host species for *Rv* highlights an overlooked factor potentially shaping biodiversity-disease relationships.

## Introduction

Wild animal populations get exposed to an array of parasite species over time, and novel parasite emergences are likely to continue with global environmental change [1–4]. The outcomes of such emergences will be shaped in part by how different co-circulating parasite species interact [5]. When multiple parasite species exploit the same host species, they may reduce [6] or increase [7] each other’s infection establishment, growth, and persistence [5,8]. Because parasites rarely emerge simultaneously [9], the sequential nature of parasite emergence can affect the strength and direction of parasite-parasite interactions and, in turn, infection prevalence in the host population [6,9,10]. Parasite interactions under sequential emergence is encapsulated in priority effects theory, and substantial evidence for priority effects in host-parasite systems indicates that timing differences in parasite emergence matter to infection outcomes [6,7,11]. Yet, rarely has the priority effects literature incorporated interactions between different host species [12], despite their known importance to infection dynamics.

Most parasite species infect multiple host species that differ in their capacity to contract, maintain and transmit infections [5,13,14]. Interspecific differences in these basic infection parameters mean that different host species can play distinct roles in parasite establishment, growth and persistence [13,14], with some host species inhibiting parasites at the host community level, and others facilitating them. The probability of parasite emergence, maintenance and proliferation can therefore differ dramatically as the relative abundance (i.e. composition) of host species in the community changes [13–15]. In particular, when two parasites have different primary hosts - those species which facilitate their establishment and persistence - there may be interactions between priority effects and multi-host community effects, whereby the strength of priority effects may be affected by host species composition [12]. In that case, any competitive advantages of emerging first, and the relative transmission and persistence rates following emergence – key factors in priority effects - will depend on the relative abundances of each primary host. Community-level differences in primary host abundance should then alter the inhibitory or facilitative effects of parasite-parasite interactions, resulting in different infection outcomes than may be observed in single-host scenarios. These predictions have yet to be empirically tested because there has yet to be formal integration of multi-parasite priority effects and multi-host dynamics.

In recent decades, parasite emergences in amphibian species have fuelled alarming biodiversity losses [16,17]. The fungal skin parasite, *Batrachochytrium dendrobatidis* (*Bd*) and the virus complex collectively called Ranavirus (*Rv*) have been particularly devasting. *Bd* and *Rv* infect a wide range of amphibian species and can cause lethal disease that has driven catastrophic amphibian mortalities globally [17,18]. Although the relative timing of *Bd* and *Rv* emergences has varied geographically and usually are not known, they were most likely asynchronous [19] and therefore potentially subject to priority effects. Notwithstanding, investigations of *Bd-Rv* interactions have produced mixed results [reviewed by 19]. Field surveillance and experiments have confirmed the occurrence of *Bd-Rv* co-infections [19,21,22], positive associations between *Bd* and *Rv* prevalence in the wild [23] and facilitative effects of prior *Bd* exposure on *Rv* success and vice versa [24]. Yet, there are also reports of negligible associations between *Bd* and *Rv* [25–27] and even negative associations [20]. In general, interactions between *Bd* and *Rv* remain poorly understood [20,22], and there is a particular knowledge gap concerning the role of *Bd*-*Rv* interactions in shaping infection dynamics in multi-host scenarios that characterize many, if not most, wild amphibian communities. Because infection prevalence following *Bd* or *Rv* emergence should increase with the relative abundance of primary host species, we may predict that *Bd* and *Rv* prevalence following sequential emergence are shaped by the interaction between the abundance of primary host species and the presence of the other parasite.

Here we experimentally assessed the extent to which the prevalence of *Bd* and *Rv* infection were influenced by host species composition, the asynchronous emergence of the other parasite, or a combination of both. We achieved these aims through a mesocosm experiment varying the presence of *Bd* and *Rv* in single- and two-host communities. We used larval common toads (*Bufo bufo*, hereafter referred to as ‘toads’) and larval common frogs (*Rana temporaria*, hereafter referred to as ‘frogs’) as focal hosts. Both species are competent hosts of *Bd* and *Rv*, but previous work suggests that toads and frogs have opposing roles in the maintenance of the two parasites [28,29]; toads are thought to be the primary host of *Bd* infection relative to frogs [30], whereas frogs appear to be the primary host of *Rv* relative to toads [28,31]. This dichotomy offered amenable conditions for examining patterns of infection proliferation following sequential *Bd-Rv* exposure in communities of multiple host species that potentially play different epidemiological roles for the two parasites.

## Methods

### Ethics statement

Approval for the animal experiment as described below, was granted by the Zoological Society of London’s Animal Welfare and Ethics Review Board (AWERB), with. procedural pathogen exposure and non-schedule 1 methods of euthanasia, permissible under the Animals in Scientific Procedures Act 1986 (United Kingdom) (project licence P8897246A).

### Animal collection and rearing

With landowner permission, we collected toad and frog egg masses from ponds near London and transported them by car to Home Office facilities at the Institute of Zoology (IoZ), Zoological Society of London (London, UK), where after a period of acclimatisation, species-specific spawn was maintained under ZSL’s standard operating procedures (SOP) and left to hatch into larvae (i.e. tadpoles). We co-housed tadpoles in plastic containers with air lines, filters, and natural plant material. The containers contained aged tap water that was tested weekly for nitrates, nitrites, and ammonia levels. Water changes of ten percent total volume were carried out every other day, and animals were fed tubifex tablets. We reared tadpoles to Gosner stage 25, at which point we transferred them to 19L plastic containers containing 15L aged tap water, co-housed by species at at stocking densities of one hundred individuals [32].

### Experimental procedures

We carried out a mesocosm experiment following a factorial design that tested the effects of sequential *Bd-Rv* exposure on the prevalence and intensity (mean infection loads) of *Bd* and *Rv* infection in both single-host and two-host communities (Fig. 1). We only considered scenarios in which *Bd* emerged before *Rv*, to reflect known emergence histories of the two parasites in Europe [19] and to keep sample sizes feasible. The experiment followed three procedures: (1) *Bd* exposure, (2) assignment of host communities, and (3) *Rv* exposure (Fig.1). For *Bd* exposures, we randomly assigned toad and frog tadpoles to one of two treatments:(1) sham exposures to sterile media (control) or (2) exposure to live *Bd* zoospores. We co-housed tadpoles in groups of 100 according to species and *Bd* treatment for the exposure period, and groups were held in 19L plastic containers containing 15L aged tap water. We administered six *Bd* doses into containers over two weeks of the experiment by pipetting 150,000 live zoospores in liquid media into the containers [33]. For sham exposures, we pipetted liquid media (lacking zoospores) into tanks at the same concentration as for *Bd* exposures.

**Fig. 1.**
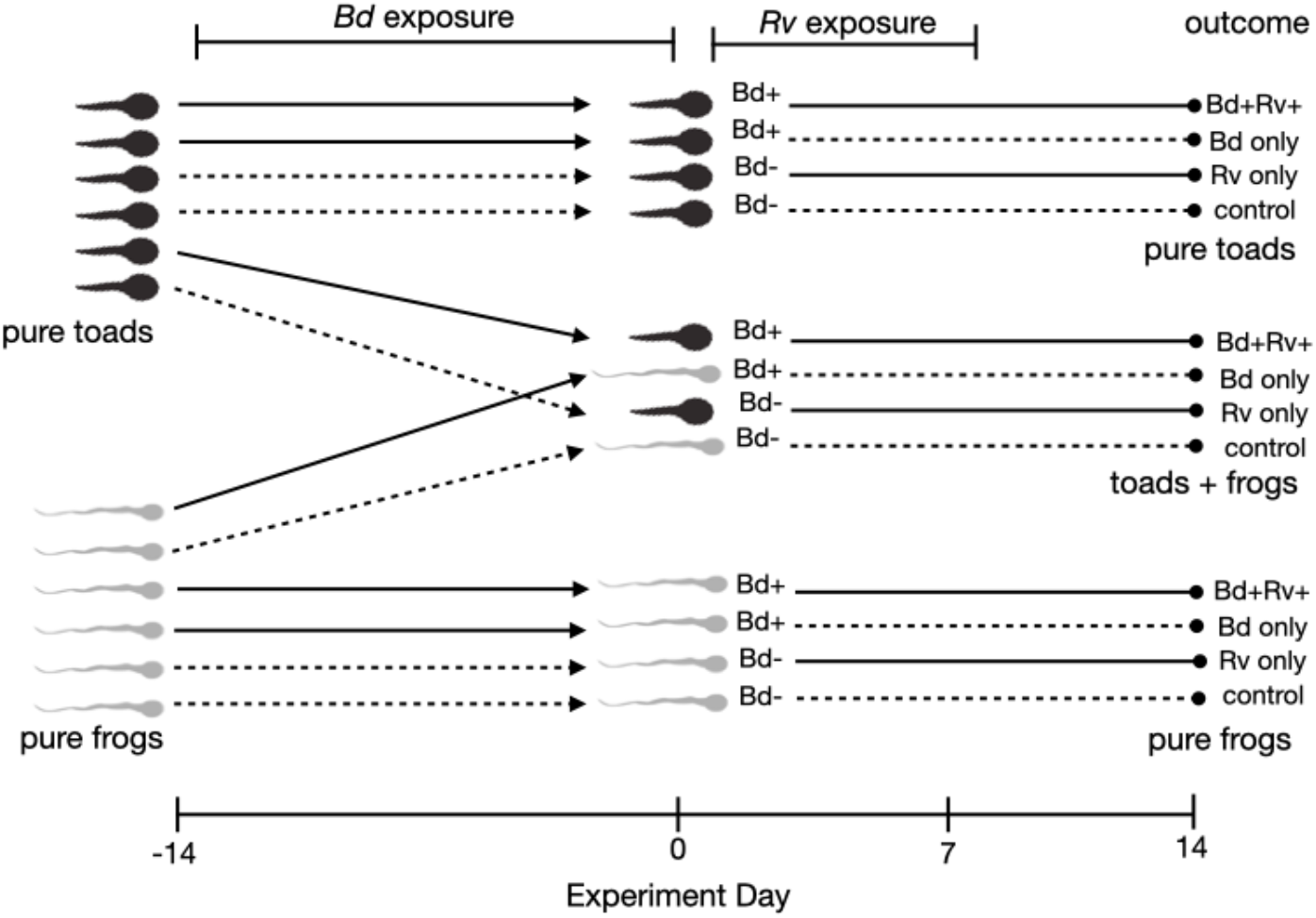
The experimental design. Toad and frog tadpoles were first co-housed separately according to their randomly assigned *Bd* exposure treatment (exposed – solid lines or unexposed – dashed lines). After two weeks of *Bd* exposure, toads and frogs were randomly assigned to one of three host community compositions (pure toads, toads + frogs, pure frogs), with total densities of ten individuals in each community. We then placed into tanks frog carcasses that were previously exposed to *Rv* (solid lines) or a sham treatment (dashed lines) to mimic subsequent *Rv* emergence. Carcasses were left in for seven days or until they were fully eaten. Day 0 of the experiment was designated as the day after *Bd* exposures, when host communities were formed and frog carcasses were placed into tanks. The experiment ended and parasite detection assays were performed 14 days later. Each tank contained 10 individuals, and each treatment combination was replicated 6 times. *Bd* = *Batrachochytrium dendrobatidis, Rv =* Ranavirus.

One week following the last *Bd* exposure, we assigned tadpoles to one of three community compositions: Pure toad communities (10 toads, 0 frogs), Pure frog communities (0 toads, 10 frogs), or mixed communities (5 toads, 5 frogs) split by *Bd* exposure treatment. We housed communities in 5L plastic containers for the duration of the experiment. Two days after assembling host communities, we assigned them to one of two *Rv* exposure scenarios: exposure to *Rv* or a sham exposure. We used the presence of *Rv*-exposed carcasses as the mode of *Rv* exposure and transmission. Specifically, we added tissue from euthanised common frog tadpoles that were previously bath exposed to either one dose of *Rv* in 14mL (10^7^ titre) in media (*Rv* present) or one dose of media only as a control (*Rv* absent). To confirm that exposures led to *Rv* infection, five reference frog tadpoles from the *Rv*-exposed group were assayed for *Rv* infection (see *Rv* assay details below). All five tested *Rv*-positive. The *Rv*-exposed and sham-exposed frog tadpoles were euthanized via flash freezing in liquid nitrogen, which also prevented decay and was a permitted non-schedule 1 method of humane euthanasia by the UK Home office. The carcasses were kept frozen until immediately before placement into experimental mesocosms, when they were thawed to room temperature. We removed carcasses from mesocosms after one week if they were not eaten.

The above procedures resulted in four parasite emergence scenarios (sham-sham, *Bd*-sham, sham-*Rv, Bd*-*Rv*), three host species compositions (pure toads, pure frogs, mixed) and twelve total treatments (2 *Bd* exposure treatments x 3 host species compositions x 2 *Rv* exposure treatments) in a fully factorial design. We replicated treatments six times, for a total of 72 experimental communities and 720 tadpoles (360 toad tadpoles, 360 frog tadpoles) (Table S1). We fed tadpoles ground tubifex tablets every other day and did ten percent water changes with cleaning on non-feeding days for the duration of the experiment. Rooms were kept at 18° Celsius with a 14-hr light cycle (6am-8pm). If humane endpoints were observed (e.g., pathological symptoms of chytridiomycosis, Ranavirosis, including and inability to self-right, feed, maintain buoyancy) euthanasia was actioned via a non-schedule 1 method (MS-222 followed by submergent in a fixative).

### Parasite detection assays

We detected *Bd* infection in tadpoles using standard molecular diagnostics involving DNA extraction and qPCR processing [34]. We extracted DNA samples from the mouthparts tadpoles, which is the only keratinized part of larval bodies that supports *Bd* infection.

Extraction of DNA was achieved using a bead-beating protocol [35], and extracts were prepared in a 1/10 dilution for qPCR. We used qPCR targeting the ITS1 gene region to measure the amount of *Bd* DNA in samples, and we ran samples in duplicate. The qPCR included negative controls and four concentration standards (0.1, 1, 10, 100) serving as positive controls [30,36,37]. We re-ran duplicate samples if only one sample amplified. *Bd* infection load is reported in genomic equivalents (GE), where one GE equals one *Bd* zoospore. A sample tested *Bd* positive when they amplified in duplicate or showed clear infection when re-running samples, and when the average GE of the two samples was greater than 0.01.

DNA was extracted from the remaining frog tadpole carcasses for *Rv* detection. Again, we used a qPCR assay, this one targeting the viral major capsid protein (MCP) gene following [38]. We included 10 standards ranging from 3 x 10^9^ – 3 viral copies to serve as positive controls. Again, we processed samples in duplicate and reran samples when only one replicate tested *Rv*-positive.

### Data Analysis

We used a generalized linear modelling (GLMM) approach in R [39] to test the effects of host species identity (toads or frogs), *Bd* and *Rv* presence (present/absent), and host species composition (toad-only, frog-only or mixed) on *Bd* and *Rv* infection outcomes, both in term of prevalence (proportion of hosts infected) and intensity (parasite load among infected hosts). Models of infection prevalence used a binomial error structure, and models of infection intensity used mean *Bd* loads, expressed as genomic equivalents (GE), as the response and Gaussian error structures. We log-transformed infection intensity values to achieve normalization and only used data from hosts testing positive for infection. We performed likelihood ratio tests based on the appropriate distribution (‘Chi-squared’ in the models of infection prevalence and ‘*F*’ in the case of infection intensity) to determine the significance of the fixed effects and their interactions on model performance (using the *dropterm* function in R).

We executed three analyses at different levels of detail. First, we assessed broad host species-level differences in infection outcomes (species-level prevalence and intensity),ignoring effects of alternative parasite presences and host species composition. We ran GLMMs for each parasite that included host species as a fixed effect and tank identity of communities as a random effect term.

Our second analysis assessed effects of alternative parasite presence and host species composition on community-level infection outcomes (community-level prevalence and intensity) without distinguishing between host species. We ran generalized linear models (GLMs), as we did not have enough power for individual-level analyses using a mixed modelling approach. The GLMs included presence of the alternative parasite, host species composition, and their interaction as fixed effects. In models of *Bd* infection prevalence and intensity, we omitted data from the ‘sham-*Rv’* treatments. Likewise, for the models of *Rv* prevalence and intensity, we omitted data from the ‘*Bd*-sham’ treatments.

The third analysis assessed community-level infection outcomes separately for frogs and toads (i.e., species-level prevalence and intensity), considering effects of host species composition and the presence of the alternative parasite. This more detailed analysis allowed us to determine whether variation in *Bd* and *Rv* infection among replicate mesocosms occurred in specific host species. We followed the same modelling approach as described for the second analysis but, as we did not have the power to run GLMs that included three-way interactions between host species, host species composition, and parasite presence, we ran separate GLMs for each host species, resulting in four GLMs for infection prevalence (one for each host and parasite combination) and four GLMs for infection intensity (one for each host and parasite combination).

We also evaluated host mortality during the experiment using similar statistical approaches. First, we explored broad interspecific differences in the likelihood of individual host mortality at the experimental endpoint, ignoring potential effects of host species composition and presence of the alternative parasite. Specifically, we ran GLMMs with individual mortality status (0 = remained alive, 1 = died prior to the experimental endpoint) as a binary response variable, species identity as a fixed effect, mesocosm identity as a random effect, and a binomial error structure. Second, we assessed per-capita host mortality in communities, considering effects of host species composition and parasite presences (none, *Bd* only, *Rv* only, *Bd-Rv*). For this analysis, we ran GLMs on tank-level data, with the proportion of total individuals in each tank that died as the response variable. The models had a binomial error structure and included host species composition and the parasite exposure scenario as fixed effects.

## Results

No animals in control groups exhibited detectable infections from either *Bd* or ranavirus. For analyses of *Bd* and *Rv* infection, we omitted data from one mesocosm due to ambiguous qPCR results (ID: 2C2, host composition = two-species, parasite presence = *Bd* only) and another mesocosm that was an outlier in terms of *Rv* infection in frog tadpoles (ID: C3C, host composition = two-species, parasite presence = *Bd* + *Rv*); all five frog tadpoles in the tank tested positive for *Rv* infection, whereas no other frog tadpoles tested positive for *Rv* infection in the other five replicate tanks with the same exposure history and host species composition. We detected co-infections in only five toad tadpoles from two-species communities, a sample size which did not permit analyses of co-infections. No clear patterns in infection intensity across treatments were evident (results presented in supplementary material). The infection results therefore focus on infection prevalence data for single *Bd* and *Rv* infections in experimental communities receiving exposure to at least one parasite. We included all mesocosms, including communities serving as controls, in analysis of mortality.

### Bd infection

There were clear interspecific host differences in the likelihood of contracting *Bd* infections (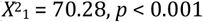 Fig. 2a). Toad tadpoles exhibited much higher prevalence of *Bd* infection at the end of the experiment (∼55%) than frog tadpoles (∼5%) irrespective of treatment (Fig. 2a). These differences persisted when scaling up to community-level *Bd* prevalence, which was strongly influenced by host composition 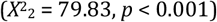. Community-level *Bd* prevalence was driven by the relative abundance of toad tadpoles; on average, *Bd* prevalence was highest in pure toad communities, lowest in pure frog communities, and intermediate in the two-host communities (Fig. 3a). We detected an interactive effect of host composition and *Rv* on *Bd* prevalence at the host community level, but the interaction was primarily driven by the strong effects of host composition. The main effect of *Rv* on community-level *Bd* prevalence was not significant 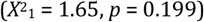.

**Fig. 2.**
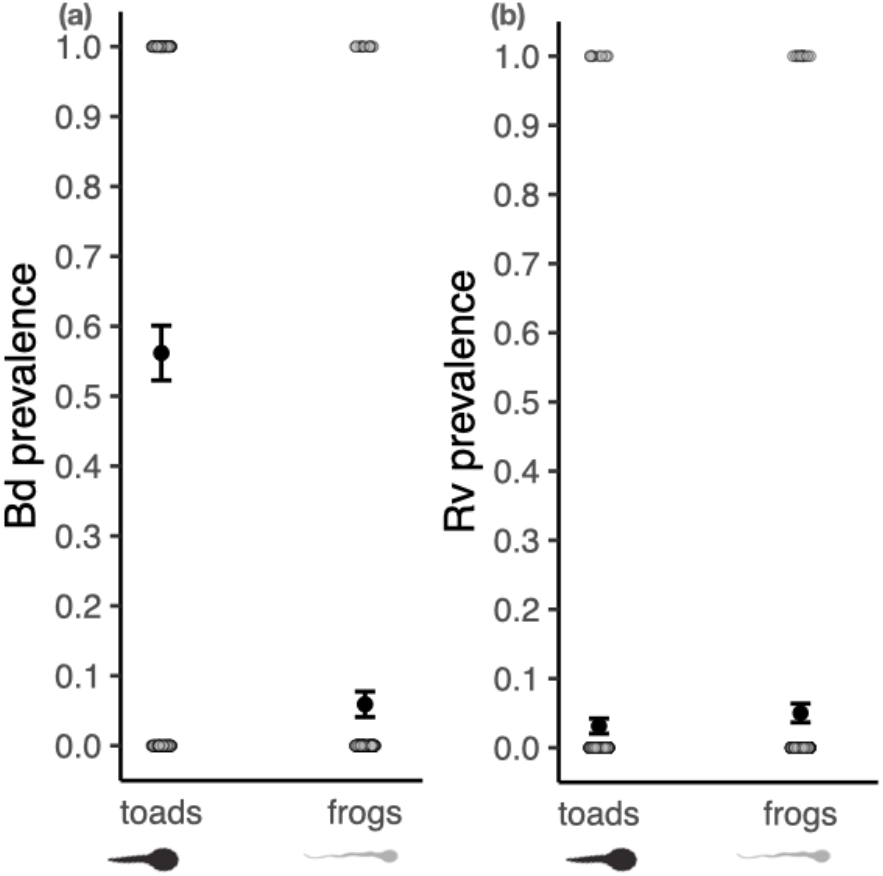
The mean prevalence of (a) *Batrachochytrium dendrobatidis (Bd)* infection and (b) Ranavirus (*Rv*) infection are shown for two focal host species: common toad tadpoles (*Bufo bufo*; ‘toads’) and common frog tadpoles (*Rana temporaria*, ‘frogs’). Infections were detected 14 days following sequential exposure to *Bd*, then *Rv*. Data are pooled across experimental communities varying in host species composition and *Bd*/Ranavirus presence. Small grey points show the raw binary values (0 = uninfected, 1 = infected). Large black points show the mean prevalence across all experimental communities. Error bars denote the standard error of the mean.

**Fig. 3.**
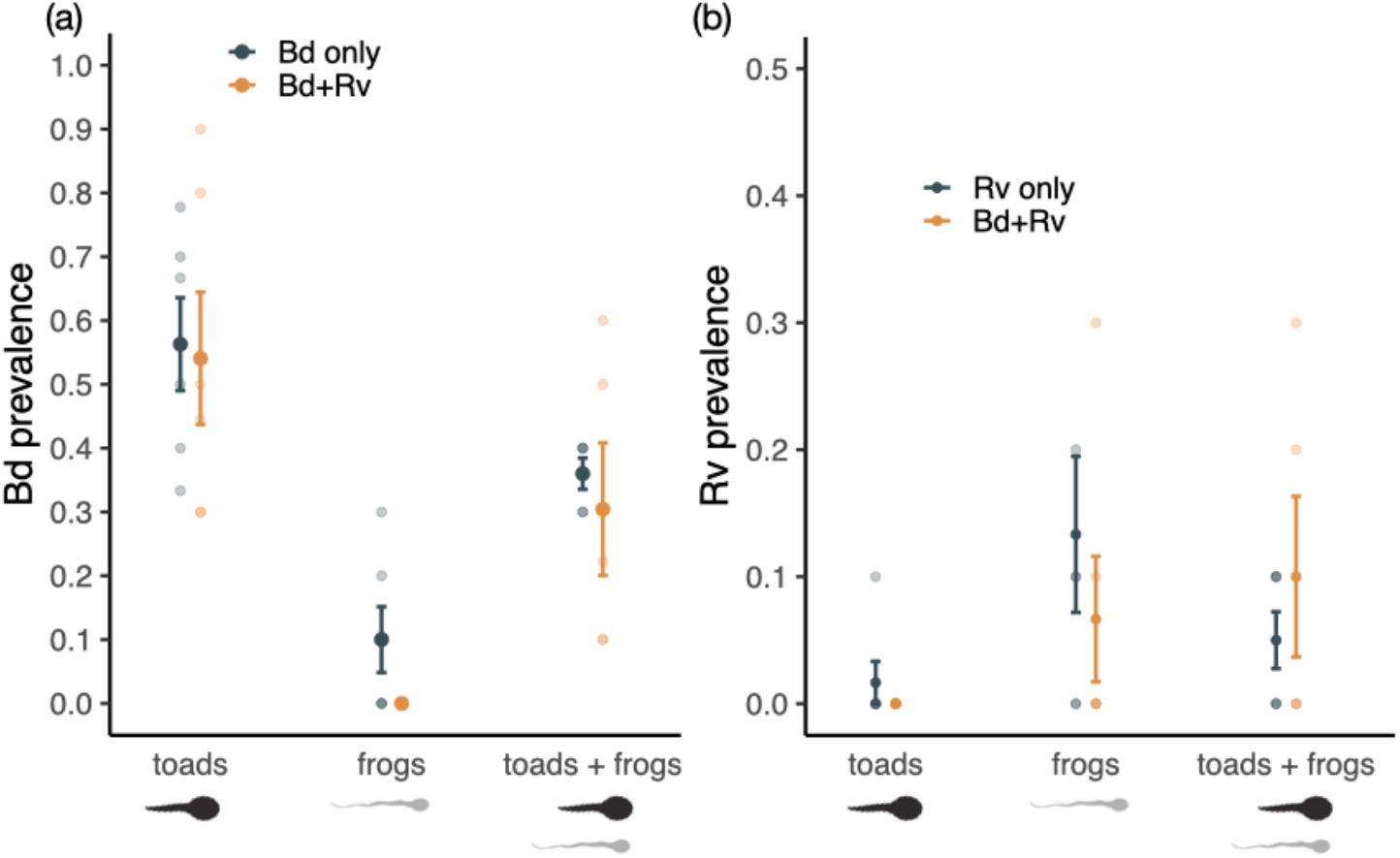
Community-level prevalence of infection of (a) *Bd* and (b) *Rv* for experimental host communities comprised of 100% toad tadpoles (toads), 100% frog tadpoles (frogs), or 50% toads and 50% frogs (toads + frogs). Each data point (small circles) represents infection prevalence in a single experimental host community. Colors denote whether communities were exposed to a single parasite (black) or both parasites (orange). Large dots denote the mean prevalence of infection among communities and error bars denote the standard error of the mean. In (b) the y-axis was rescaled to 0-50% prevalence to better illustrate different among groups. *Bd* = *Batrachochytrium dendrobatidis, Rv =* Ranavirus.

Analysing community-level *Bd* prevalence separately for frog and toad tadpoles further confirmed strong interspecific differences in *Bd* infections (Fig. 4a, b). *Bd* prevalence was consistently higher in toad tadpoles compared to frog tadpoles, irrespective of *Rv* presence and host composition (interactive effects not significant: 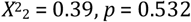 main host composition effect not significant: 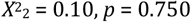 main effect of *Rv* presence not significant: (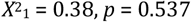 Fig. 4a). Although there were detectable interactive effects of host composition and *Rv* on *Bd* infections in frog tadpoles 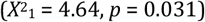, the effects were largely due to weak effects of *Rv* on *Bd* prevalence in pure frog communities (main effect of *Rv*: 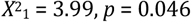 main effect of host composition: 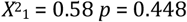 Fig. 4b); frog tadpoles subsequently exposed to *Rv* did not exhibit detectable *Bd* infections by the end of the experiment (Fig. 4b), whereas a low number of frog tadpoles in pure frog communities (∼10%) contracted *Bd* infections when not subsequently exposed to *Rv*. Despite these differences, *Bd* prevalence in frog tadpoles was generally low compared to *Bd* prevalence in toad tadpoles (Fig. 4).

**Fig. 4.**
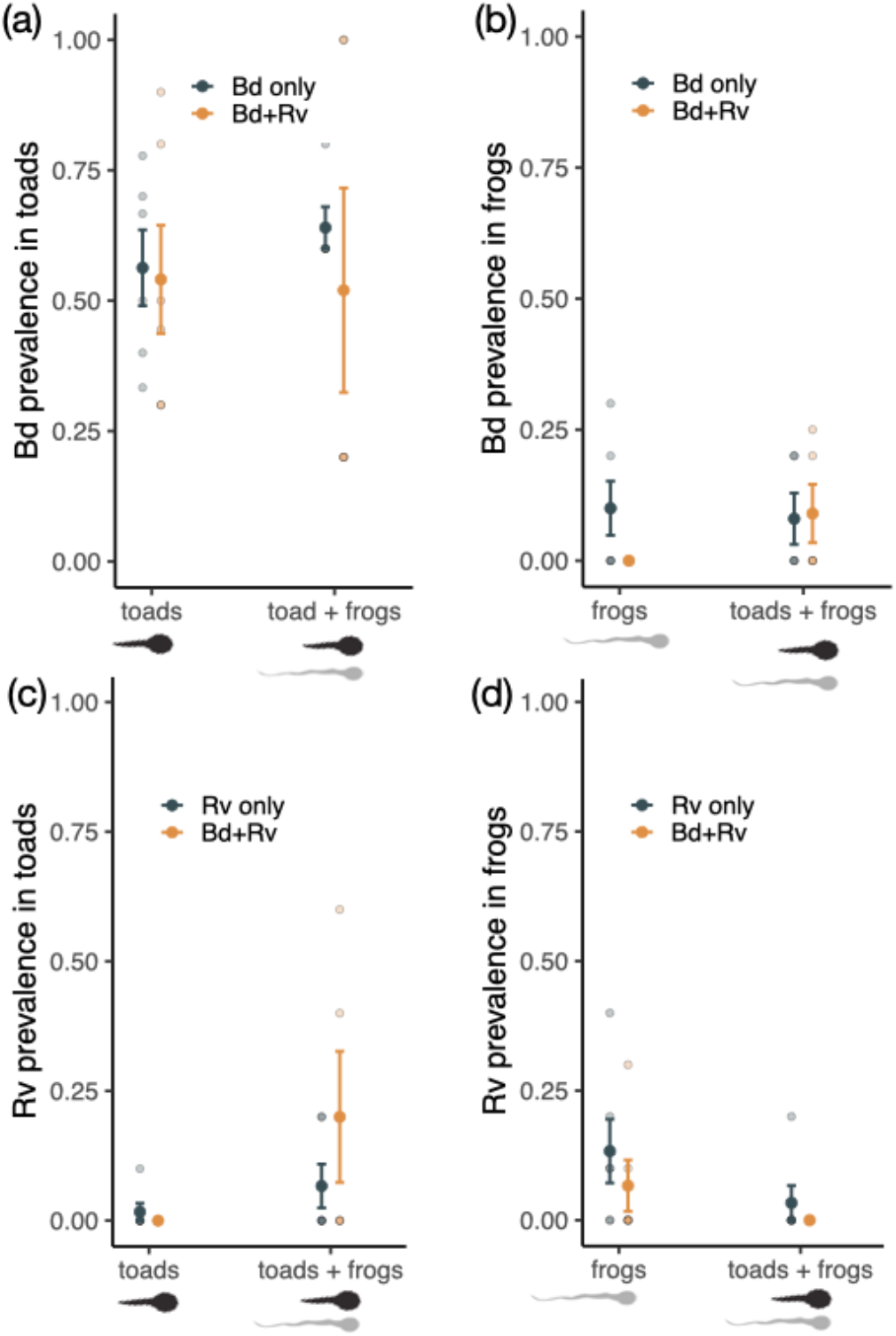
The community-level prevalence of infection of (a,b) *Bd* and (c,d) *Rv*, distinguishing by host species identity (toads – a,c, frogs – b,d). Experimental community replicates are represented by small dots, and the mean prevalence of infection in the communities is denoted by the large dots. Error bars denote the standard error of the mean prevalence of infection, and colors distinguish the *Bd* and *Rv* exposure history of the host communities. In panels for *Bd* prevalence (a,b), host communities not exposed to *Bd* were omitted, and likewise in panels for *Rv* prevalence (c,d), host communities not exposed to *Rv* were omitted. *Bd* = *Batrachochytrium dendrobatidis, Rv =* Ranavirus.

### Rv infection

*Rv* infection prevalence was generally low (∼5%) and did not broadly differ between frog tadpoles and toad tadpoles (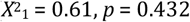 Fig. 2b). Host composition influenced community-level *Rv* prevalence: 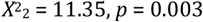), but the presence of *Bd* did not (interaction not significant: 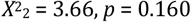 *Bd* presence: 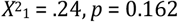 Fig. 3b).

Specifically, *Rv* prevalence was lower in toad-only communities compared to frog-only communities (GLM factor level coefficients: z < 2.41, p < 0.041). Analysing community-level *Rv* prevalence separately for frog tadpoles and toad tadpoles showed that the effects of host composition manifested in both host species, but in opposite directions (toads: 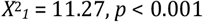 *Bd* presence not influential: 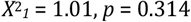, host composition x *Bd* presence interaction not influential: 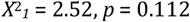 Fig. 4c. Frogs: host composition: 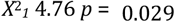, *Bd* presence not influential: 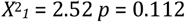, interaction not influential: 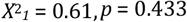 Fig. 4d). Toad tadpoles were most likely to contract *Rv* infections in two-species communities (Fig. 4c), whereas frog tadpoles were least likely to contract *Rv* infections in two-hostj communities (Fig. 4d).

### Host mortality

Forty-seven tadpoles died during the experiment (N toads = 32, N frogs = 9), two of which came from host communities serving as control groups (both were toads). Across all communities, toad tadpoles were more likely to die during the experiment than frog tadpoles 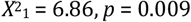 Figs. S4). At the community-level, the likelihood of mortality was influenced by host composition and the presence of either parasite (host composition: 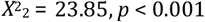 parasite presence: 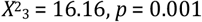 interactions were not significant: 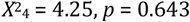 Fig. S4). Comparing the coefficients of factor levels for host composition showed that pure frog communities and two-host communities experienced fewer mortalities than pure toad communities (*z* < -3.30, *p* < 0.001; Fig. S4), and communities exposed to *Bd, Rv*, or both parasites experienced more mortality than the ‘control’ communities not exposed to either parasite (*z* >2.67, *p* < 0.008; Fig. S4)

## Discussion

Multiple generalist parasites have recently emerged in amphibian communities globally, contributing to the taxon’s imperilled status and calling for community ecological approaches to research and management. Integrating community ecological concepts on multi-parasite priority effects and multi-host community effects, our experiments showed that outcomes of sequential *Bd* and *Rv* exposure depended predominantly on host species composition, more so than priority effects arising from potential *Bd-Rv* interactions. In particular, infection outcomes in single-host communities did not mirror infection outcomes in two-host communities for either parasite. Furthermore, the mechanisms driving different infection outcomes in multi-host communities were specific to each parasite.

Infection dynamics in multi-host systems can be altered by host species composition simply through changes in the abundance of primary host species [13,15,40,41], forming a basis for biodiversity-disease relationships [15,40,41]. This mechanism largely explained the effect of host species composition on *Bd* infection. Regardless of *Rv* presence, toad tadpoles were much more likely to contract *Bd* infections than frog tadpoles, consistent with previous work [37,42]. Furthermore, since toad tadpoles were consistently the apparent primary host of *Bd*, community-level *Bd* prevalence declined with decreasing abundance of toad tadpoles in communities (i.e., moving from the pure toad communities with ten toads to the mixed communities with five toads). Notably, the number of toad deaths among communities also decreased when going from *Bd*-exposed pure toad communities to *Bd*-exposed two-host communities, suggesting that *Bd* infections in toad tadpoles may have driven patterns of host mortality. This is unusual, in that most research has supported that conclusion that mortality associated with chytridiomycosis in anurans is restricted to post-metamorphic stages [16].

However, low levels of mortality have been reported in common toad tadpoles, presumably due to indirect costs of infection [37].

We saw different patterns when examining *Rv* dynamics. Going from single-host to two-host communities changed not only the overall, community-level *Rv* prevalence but also *Rv* infection probabilities in both host species. Strikingly, the *Rv* infection probabilities in frog and toad tadpoles changed in opposing directions when moving from single-to two-host communities; co-occurrence of the two host species increased the likelihood of *Rv* infection in toad tadpoles but decreased the likelihood of *Rv* infection in frog tadpoles, essentially reversing the apparent primary host for *Rv* between single-vs. two-host communities. This reversal highlights that the species serving as the apparent primary hosts for a given parasite may be dynamic, changing with host community composition and relative abundances of the different host species. As such, species-specific variation in infection probability may obscure predictions for biodiversity-disease relationships, which generally assume fixed infection probabilities across different host community compositions [41,43].

Determining the transmission dynamics of *Rv* in multi-host scenarios may reveal explanations for the surprising reversals in the parasite’s apparent primary host species. There were two possible routes of *Rv* transmission in the mesocosms: direct, via host-to-host transmission, and indirect via consumption of infected carcass. Hence multiple scenarios involving these two transmission routes may have generated the observed reversals in the apparent primary host species. We propose two hypotheses: an indirect transmission hypothesis and a direct transmission hypothesis. Under the indirect transmission hypothesis, interspecific differences in indirect transmission via feeding on dead carcasses explained the reversals. For example, toad tadpoles may have become more aggressive feeders when frog tadpoles were present, similar to context-dependent behaviours observed in competing fish species [44], and hence became more exposed to *Rv* infective stages on the carcasses (increasing prevalence in toads), and potentially reducing exposure for frogs (decreasing prevalence in frogs). Conversely, the direct transmission hypothesis assumes that frog tadpoles are more infectious for *Rv* than toad tadpoles, as has been argued previously [28,45], and act as the primary source of all *Rv* infections in the community. Hence, the two-host communities would have brought toads into contact with more highly infectious hosts (frog tadpoles) than in pure toad communities (five potentially highly infectious frog tadpoles in two-host communities versus zero in pure toad communities), increasing prevalence in toads. The opposite would also have been true for frog tadpoles (up to ten potentially highly infectious frog tadpoles in one-host communities vs. only five in the two-host communities), reducing prevalence in frogs. Furthermore, while this effect purely relies on the relative abundance of highly infectious frogs and low infectious toads, the effect may be exacerbated if low-competent toad tadpoles remove viral infective stages from the environment, reducing onward transmission opportunities to frogs via a transmission interference dilution effect [40,43]. This dilution effect was posited by North et al. (2015) after observing in wild frog populations (*R. temporaria*) that *Rv* prevalence was negatively associated with abundance of co-occurring wild toads (*B. bufo*). Despite these coarse lines of evidence, all the above hypotheses remain speculative due to a lack of knowledge on species-specific *Rv* shedding and resulting transmission patterns that determine host competence.

As one of the first formal comparisons of sequential infection dynamics in single-versus multi-host communities, this study demonstrates the insights afforded through the integration of multi-parasite priority effects with multi-host infection dynamics. Two main insights emerged for *Bd*-*Rv* dynamics. First, the patterns of *Bd* and *Rv* infection in single-host contexts were not representative of those we observed in multi-host contexts, meaning that, at least in the case of *Bd* and *Rv*, sequential infection dynamics in single-host systems may have limited applicability to multi-host amphibian communities common in the wild. Second and most crucially, patterns of *Bd* and *Rv* infection following sequential emergence were shaped predominately by multi-host effects, not multi-parasite effects. Despite creating the potential for interaction between *Bd* and *Rv* in our experiments, any such interactions were weak at best and did little to moderate the unambiguous influence of host species composition on *Bd* and *Rv* infection. In terms of *Bd* and *Rv* risk, multi-host effects appear to dominate over multi-parasite effects, suggesting that efforts to manage biodiversity threats from the two parasites should focus interventions on host community assemblages rather than controlling *Bd* and *Rv* co-occurrence.

## References

1. Altizer S, Ostfeld RS, Johnson PTJ, Kutz S, Harvell CD. 2013 Climate change and infectious diseases: From evidence to a predictive framework. Science 341, 514–519. (doi:10.1126/science.1239401)

2. Mahon MB et al. 2024 A meta-analysis on global change drivers and the risk of infectious disease. Nature (doi:10.1038/s41586-024-07380-6)

3. Cohen JM, Sauer EL, Santiago O, Spencer S, Rohr JR. 2020 Divergent impacts of warming weather on wildlife disease risk across climates. Science 370, eabb1702. (doi:10.1126/science.abb1702)

4. Gallana M, Ryser-Degiorgis M-P, Wahli T, Segner H. 2013 Climate change and infectious diseases of wildlife: Altered interactions between pathogens, vectors and hosts. Current Zoology 59, 427–437. (doi:10.1093/czoolo/59.3.427)

5. Johnson PTJ, de Roode JC, Fenton A. 2015 Why infectious disease research needs community ecology. Science 349, 1259504–1259504. (doi:10.1126/science.1259504)

6. de Roode JC, Helinski MEH, Anwar MA, Read AF. 2005 Dynamics of multiple infection and within-host competition in genetically diverse malaria infections. The American Naturalist 166, 531–542.

7. Halliday FW, Penczykowski RM, Barrès B, Eck JL, Numminen E, Laine A-L. 2020 Facilitative priority effects drive parasite assembly under coinfection. Nat Ecol Evol 4, 1510–1521. (doi:10.1038/s41559-020-01289-9)

8. Rynkiewicz EC, Pedersen AB, Fenton A. 2015 An ecosystem approach to understanding and managing within-host parasite community dynamics. Trends in Parasitology 31, 212–221. (doi:10.1016/j.pt.2015.02.005)

9. Karvonen A, Jokela J, Laine A-L. 2019 Importance of sequence and timing in parasite coinfections. Trends in Parasitology 35, 109–118. (doi:10.1016/j.pt.2018.11.007)

10. Clay PA, Cortez MH, Duffy MA, Rudolf VHW. 2019 Priority effects within coinfected hosts can drive unexpected population-scale patterns of parasite prevalence. Oikos 128, 571–583. (doi:10.1111/oik.05937)

11. Clay PA, Duffy MA, Rudolf VHW. 2020 Within-host priority effects and epidemic timing determine outbreak severity in co-infected populations. Proc. R. Soc. B. 287, 20200046. (doi:10.1098/rspb.2020.0046)

12. Jiao J, Cortez MH. 2022 Exploring how a generalist pathogen and within-host priority effects alter the risk of being infected by a specialist pathogen. The American Naturalist 200, 815–833. (doi:10.1086/721762)

13. Daversa D, Bosch J, Manica A, Garner TW, Fenton A. 2022 Host identity matters – up to a point: the community context of Batrachochytrium dendrobatidis transmission. The American Naturalist, 720638. (doi:10.1086/720638)

14. Fenton A, Streicker DG, Petchey OL, Pedersen AB. 2015 Are all hosts created equal? Partitioning host species contributions to parasite persistence in multihost communities. The American Naturalist 186, 610–622. (doi:10.1086/683173)

15. Johnson PTJ, Preston DL, Hoverman JT, Richgels KLD. 2013 Biodiversity decreases disease through predictable changes in host community competence. Nature 494, 230–233. (doi:10.1038/nature11883)

16. Fisher MC, Garner TWJ. 2020 Chytrid fungi and global amphibian declines. Nat Rev Microbiol 18, 332–343. (doi:10.1038/s41579-020-0335-x)

17. Price SJ, Garner TWJ, Nichols RA, Balloux F, Ayres C, Mora-Cabello de Alba A, Bosch J. 2014 Collapse of amphibian communities due to an introduced ranavirus. Current Biology 24, 2586–2591. (doi:10.1016/j.cub.2014.09.028)

18. Scheele BC et al. 2019 Amphibian fungal panzootic causes catastrophic and ongoing loss of biodiversity. Science 363, 1459–1463. (doi:10.1126/science.aav0379)

19. Rosa GM et al. 2017 Impact of asynchronous emergence of two lethal pathogens on amphibian assemblages. Sci Rep 7, 43260. (doi:10.1038/srep43260)

20. Herczeg D, Ujszegi J, Kásler A, Holly D, Hettyey A. 2021 Host–multiparasite interactions in amphibians: a review. Parasites Vectors 14, 296. (doi:10.1186/s13071-021-04796-1)

21. Souza MJ, Gray MJ, Colclough P, Miller DL. 2012 Prevalence of infection by Batrachochytrium dendrobatidis and Ranavirus in Eastern Hellbenders (Cryptobranchus alleganiensis alleganiensis) in eastern Tennessee. Journal of Wildlife Diseases 48, 560–566. (doi:10.7589/0090-3558-48.3.560)

22. Warne RW, LaBumbard B, LaGrange S, Vredenburg VT, Catenazzi A. 2016 Co-Infection by chytrid fungus and ranaviruses in wild and harvested frogs in the tropical Andes. PLoS ONE 11, e0145864. (doi:10.1371/journal.pone.0145864)

23. Stutz WE, Blaustein AR, Briggs CJ, Hoverman JT, Rohr JR, Johnson PTJ. 2018 Using multi-response models to investigate pathogen coinfections across scales: Insights from emerging diseases of amphibians. Methods Ecol Evol 9, 1109–1120. (doi:10.1111/2041-210X.12938)

24. Ramsay C, Rohr JR. 2021 The application of community ecology theory to co-infections in wildlife hosts. Ecology 102, e03253. (doi:10.1002/ecy.3253)

25. Olori J, Netzband R, McKean N, Lowery J, Parsons K, Windstam S. 2018 Multi-year dynamics of ranavirus, chytridiomycosis, and co-infections in a temperate host assemblage of amphibians. Dis. Aquat. Org. 130, 187–197. (doi:10.3354/dao03260)

26. Bosch J, Monsalve-Carcaño C, Price SJ, Bielby J. 2020 Single infection with (Batrachochytrium dendrobatidis) or Ranavirus does not increase probability of co-infection in a montane community of amphibians. Sci Rep 10, 21115. (doi:10.1038/s41598-020-78196-3)

27. Herczeg D, Holly D, Kásler A, Bókony V, Papp T, Takács-Vágó H, Ujszegi J, Hettyey A. 2023 Amphibian larvae benefit from a warm environment under simultaneous threat from chytridiomycosis and ranavirosis. Oikos 2023, e09953. (doi:10.1111/oik.09953)

28. Duffus ALJ, Nichols RA, Garner TWJ. 2014 Experimental evidence in support of single host maintenance of a multihost pathogen. Ecosphere 5, art142. (doi:10.1890/ES14-00074.1)

29. Clare FC et al. 2016 Climate forcing of an emerging pathogenic fungus across a montane multi-host community. Philosophical Transactions of the Royal Society B: Biological Sciences 371, 20150454. (doi:10.1098/rstb.2015.0454)

30. Bielby J, Fisher MC, Clare FC, Rosa GM, Garner TWJ. 2015 Host species vary in infection probability, sub-lethal effects, and costs of immune response when exposed to an amphibian parasite. Scientific Reports 5, 10828. (doi:10.1038/srep10828)

31. Ford CE, Brookes LM, Skelly E, Sergeant C, Jordine T, Balloux F, Nichols RA, Garner TWJ. 2022 Non-Lethal Detection of Frog Virus 3-Like (RUK13) and Common Midwife Toad Virus-Like (PDE18) Ranaviruses in Two UK-Native Amphibian Species. Viruses 14, 2635. (doi:10.3390/v14122635)

32. Gosner KL. 1960 A simplified table for staging anuran embryos and larvae with notes on identification. Herpetologica 16.

33. Daversa DR, Manica A, Bosch J, Jolles JW, Garner TWJ. 2018 Routine habitat switching alters the likelihood and persistence of infection with a pathogenic parasite. Functional Ecology 32, 1262–1270. (doi:10.1111/1365-2435.13038)

34. Boyle DG, Boyle DB, Olsen V, Morgan JAT, Hyatt AD. 2004 Rapid quantitative detection of chytridiomycosis (Batrachochytrium dendrobatidis) in amphibian samples using real-time Taqman PCR assay. Diseases of aquatic organisms 60, 141–148.

35. Bates KA et al. 2022 Microbiome function predicts amphibian chytridiomycosis disease dynamics. Microbiome 10. (doi:10.1186/s40168-021-01215-6)

36. Luquet E, Garner TWJ, Léna J-P, Bruel C, Joly P, Lengagne T, Grolet O, Plénet S. 2012 Genetic erosion in wild populations makes resistance to a pathogen more costly. Evolution 66, 1942–1952. (doi:10.1111/j.1558-5646.2011.01570.x)

37. Garner TWJ, Walker S, Bosch J, Leech S, Marcus Rowcliffe J, Cunningham AA, Fisher MC. 2009 Life history tradeoffs influence mortality associated with the amphibian pathogen Batrachochytrium dendrobatidis. Oikos 118, 783–791. (doi:10.1111/j.1600-0706.2008.17202.x)

38. Leung WTM, Thomas-Walters L, Garner TWJ, Balloux F, Durrant C, Price SJ. 2017 A quantitative-PCR based method to estimate ranavirus viral load following normalisation by reference to an ultraconserved vertebrate target. Journal of Virological Methods 249, 147–155. (doi:10.1016/j.jviromet.2017.08.016)

39. R Core Team. 2023 R: A language and environment for statistical computing.

40. Ostfeld RS, Keesing F. 2012 Effects of host diversity on infectious disease. Annu. Rev. Ecol. Evol. Syst. 43, 157–182. (doi:10.1146/annurev-ecolsys-102710-145022)

41. Johnson PTJ, Stewart Merrill TE, Dean AD, Fenton A. 2024 Diverging effects of host density and richness across biological scales drive diversity-disease outcomes. Nat Commun 15, 1937. (doi:10.1038/s41467-024-46091-4)

42. Baláž V et al. 2014 Assessing risk and guidance on monitoring of Batrachochytrium dendrobatidis in Europe through identification of taxonomic selectivity of infection. Conservation Biology 28, 213–223. (doi:10.1111/cobi.12128)

43. Keesing F, Holt RD, Ostfeld RS. 2006 Effects of species diversity on disease risk: Effects of species diversity on disease risk. Ecology Letters 9, 485–498. (doi:10.1111/j.1461-0248.2006.00885.x)

44. Taniguchi Y, Nakano S. 2000 Condition-specific competition: Implications for the altitudinal distribution of stream fishes. Ecology 81, 2027–2039. (doi:10.1890/0012-9658(2000)081[2027:CSCIFT]2.0.CO;2)

45. North AC, Hodgson DJ, Price SJ, Griffiths AGF. 2015 Anthropogenic and Ecological Drivers of Amphibian Disease (Ranavirosis). PLoS ONE 10, e0127037. (doi:10.1371/journal.pone.0127037)

