## Supplementary Material for "Host species composition, not priority effects, determines infection risk in multihost-multiparasite amphibian communities"

| | **Host species composition** | | | | | | |
|---|---|---|---|---|---|---|---|
| **Parasite scenario** | **toads only** | | **frogs only** | | **toads + frogs** | | |
| | **toad count** | **frog count** | **toad count** | **frog count** | **Toad count** | **frog count** | |
| **sham + sham (control)** | 10 | 0 | 0 | 10 | 5 | 5 | |
| **Bd + sham (Bd only)** | 10 | 0 | 0 | 10 | 5 | 5 | |
| ***Bd* + *Rv*** | 10 | 0 | 0 | 10 | 5 | 5 | |
| **sham + *Rv* (*Rv* only)** | 10 | 0 | 0 | 10 | 5 | 5 | |
| **Per treatment total** | 40 | 0 | 0 | 40 | 20 | 20 | **120** |
| **X 6 replicates** | 240 | 0 | 0 | 240 | 120 | 120 | **720** |

**Table S1. Original sample sizes for the mesocosm experiment.** The mesocosm experiment varied host species composition (columns) and parasite emergence scenario (rows), while keeping total host densities among treatments constant. The factorial design resulted in 12 treatments and were replicated six times, resulting in 720 individual hosts grouped into 72 mesocosms. ‘Toads’ is short for common toad (*Bufo bufo*), and ‘frogs’ is short for common frogs (*Rana temporaria*). All hosts were in the larval ‘tadpole’ stage at the time of the experiment. Data from two mesocosms (N = 1 Bd only, frogs + toads; 1 Bd+Rv, frogs + toads) were omitted from the analysis, one due to ambiguity in the PCR data and another due to the data being outliers.

**Data analysis of infection intensity**

We repeated the three data analyses described in the Methods using *Bd* and *Rv* infection intensity as response variables. We measured *Bd* infection intensity (i.e. *Bd* load) in genomic equivalents (GE), whereby one GE denotes a single zoospore detected on samples from qPCR assays [1]. We measured *Rv* intensity (*Rv* load) as the number of virus copies detected on samples from the qPCR assays. We considered only *Bd*- and *Rv*-positive samples for infection intensity responses, and we log10-transformed values to achieve normalization.

*Results*

We did not detect any general interspecific differences in *Bd* infection or *Rv* infections loads (*Bd*: $X^2_1$ = 0.21, $p$ = 0.640; *Rv*: $X^2_1$ = 0.59, $p$ = 0.439; Fig. S1). We did detect an interactive effect of *Rv* presence and host composition on mean *Bd* loads at the tank level ($F$ = 5.51, $p$ = 0.029), but there were no clear patterns (Fig. S2). We did not detect interactive or main effects of *Bd* presence or host composition on mean *Rv* loads at the community level (interactive effects: $F$ = 0.02, $p$ = 0.900; main effect of *Bd* presence: $F$ = 0.10, $p$ = 0.910; main effect of host composition: $F$ = 1.64, $p$ = 0.236). When breaking community-level mean infection intensities down by species, we found no main effects (we did not have the power to examine interactive effects) of *Rv* presence or host composition on mean *Bd* intensity in toad tadpoles (*Rv* presence: $F$ = 1.50, $p$ = 0.235, main effect of host composition: $F$ = 0.01, $p$ = 0.934) or frog tadpoles (*Bd* presence: $F$ = 0.15, $p$ = 0.720, host composition: $F$ = 0.51, $p$ = 0.515). We also did not detect any interactive or main effects of *Bd* presence and host composition on mean *Rv* loads in toad tadpoles (*Bd* presence: $F$ = 0.21, $p$ = 0.692, host composition: $F$ = 0.29, $p$ = 0.643) or frog tadpoles (main effect of *Bd* presence: $F$ = 0.64, $p$ = 0.469, main effect of host composition: $F$ = 0.29, $p$ = 0.616)

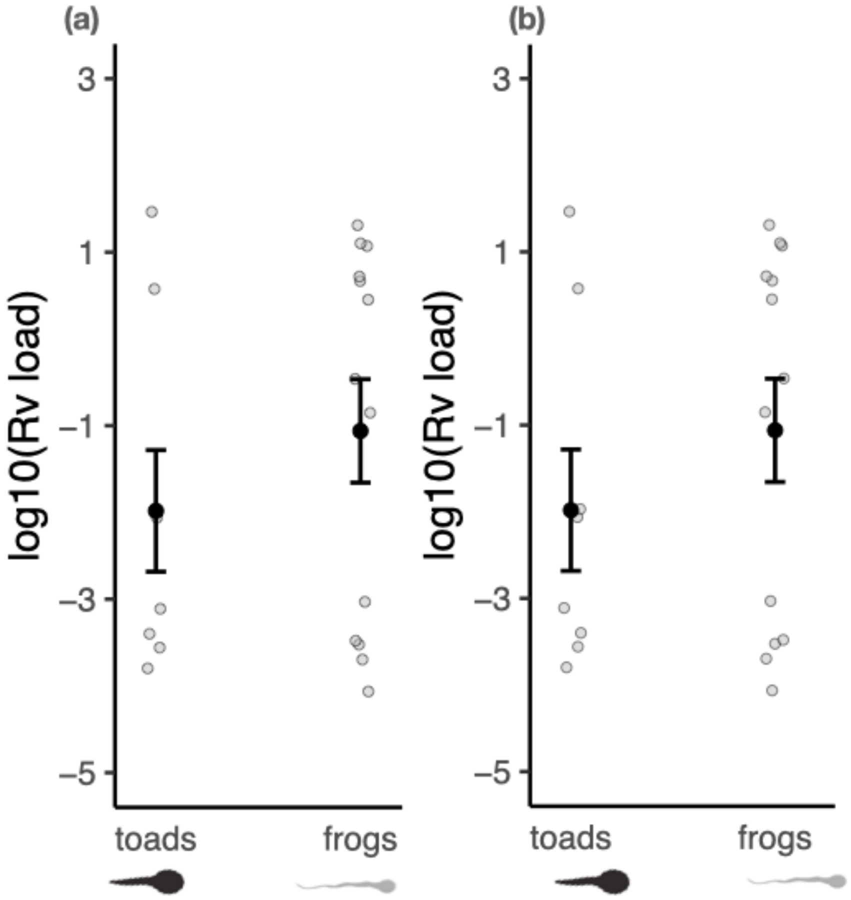


**Fig. S1.** The mean intensity of (a) *Batrachochytrium dendrobatidis (Bd)* infection and *(b)* Ranavirus (*Rv*) infection for two focal host species: common toads (*Bufo bufo*; 'toads') and common frogs (*Rana temporaria*, 'frogs'). Infection intensities are reported as log10-transformed genomic equivalents for *Bd* and log10-transformed number of viral copies, both measured through qPCR assays. Data are pooled across experimental communities varying in parasite exposure scenario and host species composition. Small grey points show the mean infection intensities for each community. Large black points show the overall mean infection intensities across all experimental communities.  Error bars denote the standard error of the mean.

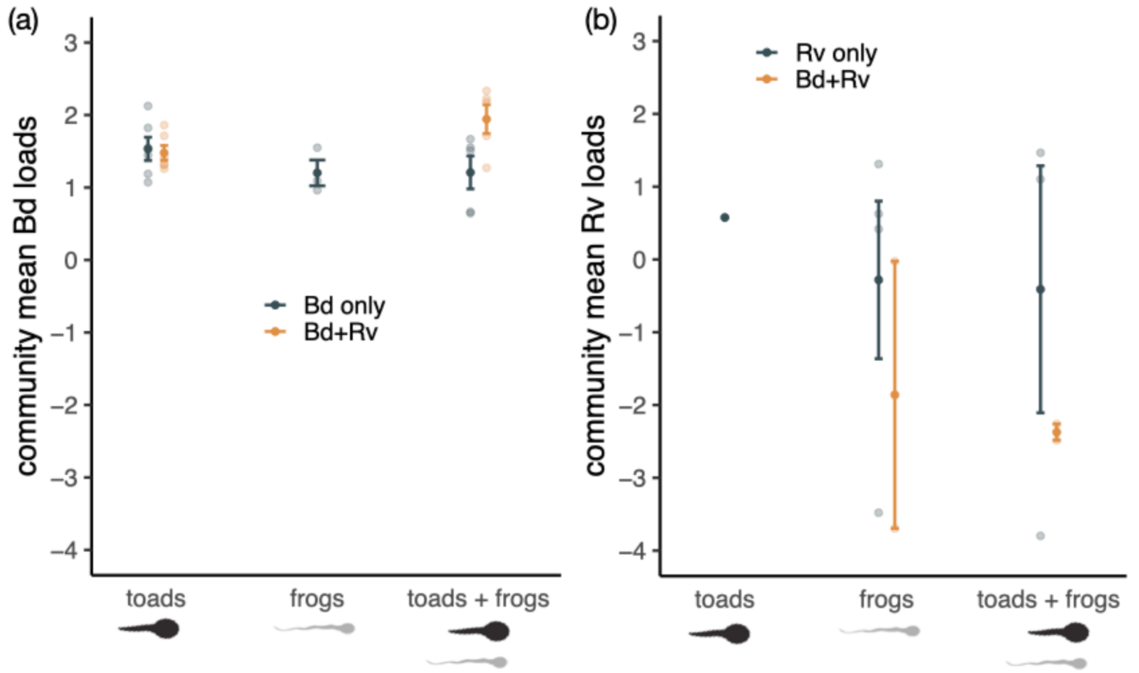


**Fig. S2.** Variation in (a) *Bd* infection intensity and (b) *Rv* infection intensity across communities varying in host species composition (x axis) and parasite emergence scenario (line color). Infection intensities are reported as log10-transformed genomic equivalents for *Bd* and log10-transformed number of viral copies, both measured through qPCR assays. Lines denote the mean infection intensity, and error bars denote the standard error of the mean. Only data from animals testing positive for infection are shown. *Bd* = *Batrachochytrium dendrobatidis, Rv* = Ranavirus*.*

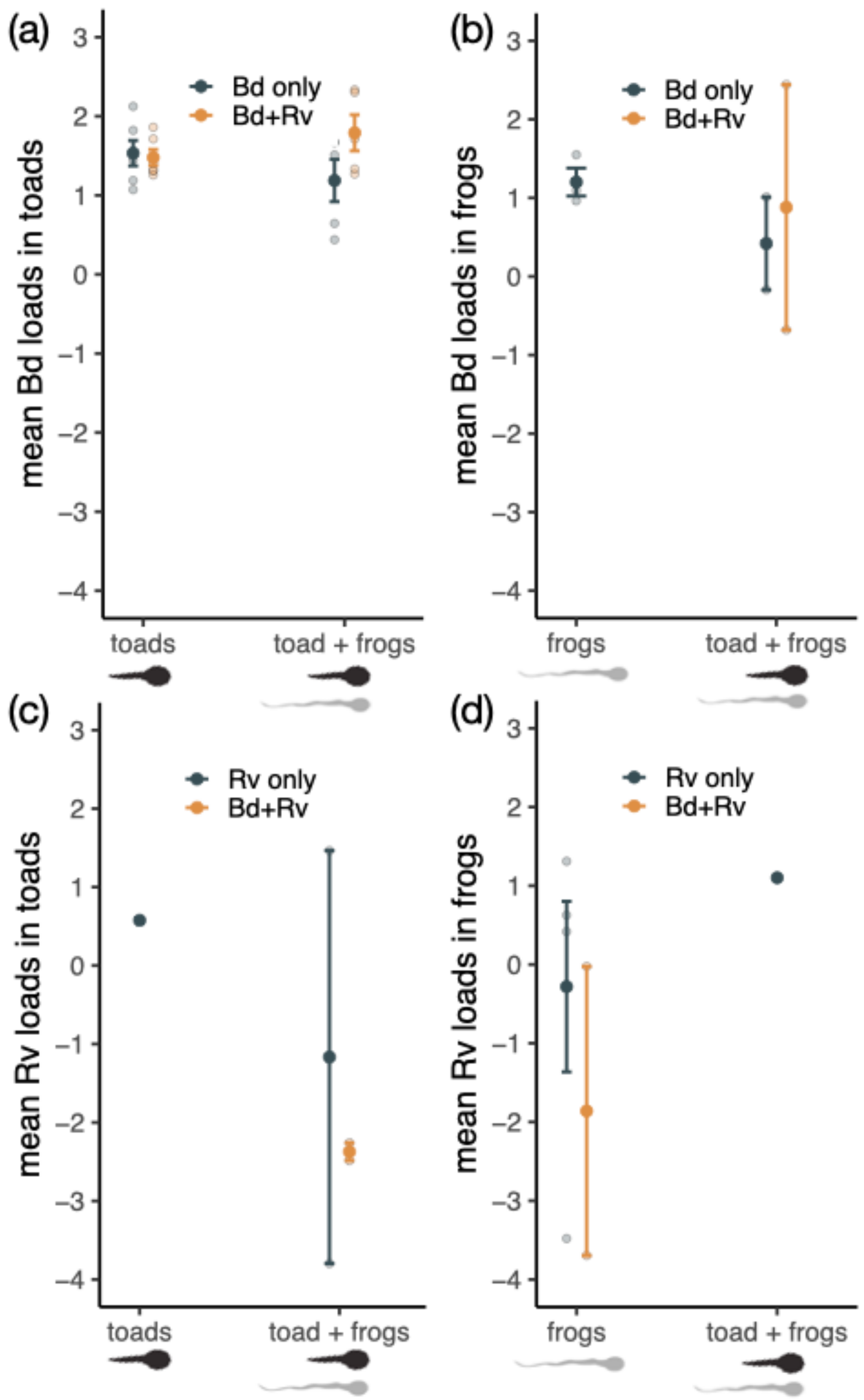


**Fig. S3.** The community-level intensities of infection from (a,b) *Bd* and (c,d) *Rv*, distinguishing host species identity (toads – a,c, frogs – b,d). Infection intensities are reported as log10-transformed genomic equivalents for *Bd* and log10-transformed number of viral copies, both measured through qPCR assays. Experimental community replicates are represented by small dots, and the mean infection intensities of communities are denoted by the large dots. Error bars denote the standard error of the mean infection intensity, and colors distinguish the *Bd* and *Rv* exposure history of the host communities. In panels for *Bd* intensity (a,b), host communities not exposed to *Bd* were omitted, and likewise in panels for *Rv* intensity(c,d), host communities not exposed to *Rv* were omitted. Only data from individuals testing positive for infection were included. *Bd* = *Batrachochytrium dendrobatidis, Rv* = Ranavirus.

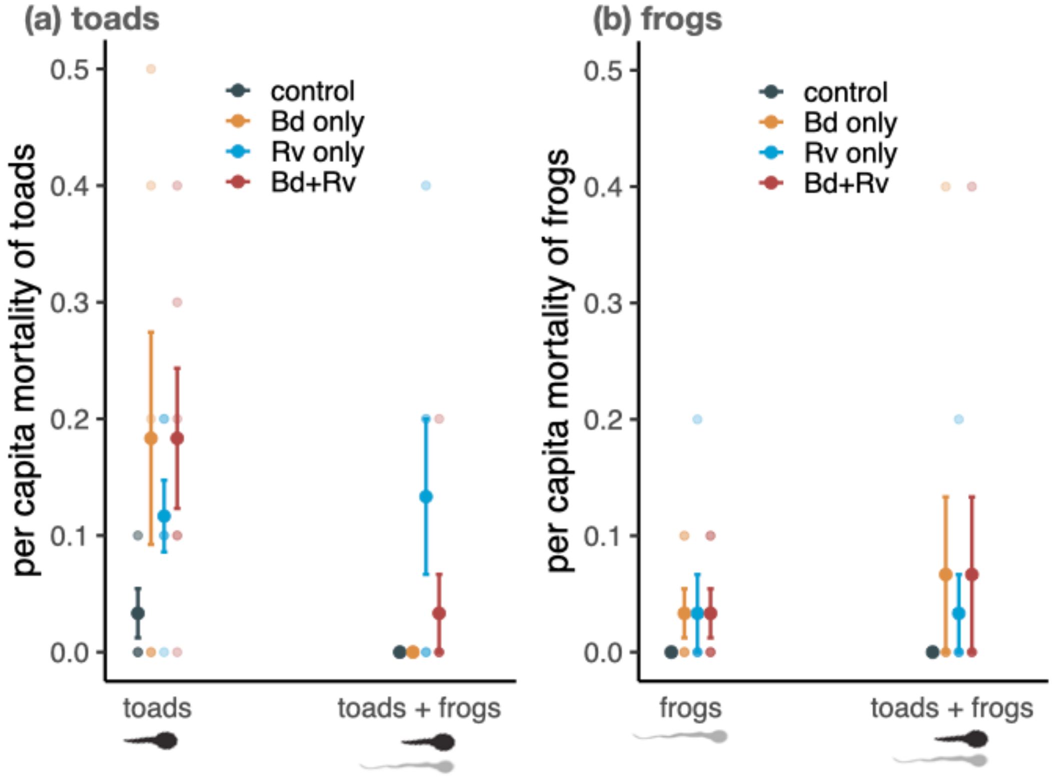


**Fig. S4.** Per-capita host mortality of experimental communities varying in host species composition (pure toads, pure frogs, toads + frogs) is displayed separately for toad tadpoles (left panel) and frog tadpoles (right panel). Points represent mean proportion of individuals that died before the end of the experiment, and error bars denote the standard error of the mena. Colors denote the parasite exposure treatment [exposure only to *Batrachochytrium dendrobatidis* (*Bd* only), exposure only to Ranavirus (*Rv* only), or sequential *Bd-Rv* exposure (*Bd + Rv*)].
